# A Real-Time Analysis of Protein Transport via the Twin Arginine Translocation Pathway in Response to Different Components of the Protonmotive Force *in vivo*

**DOI:** 10.1101/2023.01.12.523868

**Authors:** Wenjie Zhou, Binhan Hao, Terry M. Bricker, Steven M. Theg

## Abstract

The twin arginine translocation (Tat) pathway transports folded protein across the cytoplasmic membrane in bacteria, archaea, and across the thylakoid membrane in plants as well as the inner membrane in some mitochondria. In plant chloroplasts, the Tat pathway utilizes the protonmotive force (PMF) to drive protein translocation. However, in bacteria, it has been shown that Tat transport depends only on the Δψ component of PMF *in vitro*. To investigate the comprehensive PMF requirement in *Escherichia coli*, we have developed the first real-time assay to monitor Tat transport utilizing the NanoLuc Binary Technology (NanoBiT) in *E. coli* spheroplasts. This luminescence assay allows for continuous monitoring of Tat transport with high-resolution, making it possible to observe subtle changes in transport in response to different treatments. By applying the NanoLuc assay, we report that, under acidic conditions, ΔpH, in addition to Δψ, contributes energetically to Tat transport *in vivo* in *E. coli* spheroplasts. These results provide novel insight into the mechanism of energy utilization by the Tat pathway.

## Introduction

The twin arginine translocation (Tat) pathway, which is present in bacteria, archaea, thylakoids in chloroplasts, and some mitochondria, is dedicated to transporting proteins in their folded conformation across energy-transducing membranes. This pathway plays a critical role in a variety of biosynthetic pathways, including energy metabolism, heavy metal resistance, cell division, virulence in prokaryotes, as well as photosynthesis in chloroplasts (1–3). The minimal Tat translocon consists of the TatA and TatC proteins, although in most systems, TatB is also required (1, 4, 5). Transport via the Tat pathway is dependent on the protonmotive force (PMF), which is composed of the transmembrane electrical potential (Δψ) and proton gradient (ΔpH) (2, 6–8). However, how Tat protein transport is coupled energetically with to the PMF is unclear.

A detailed mechanistic understanding of the Tat transport mechanism is limited by current Tat transport assays. Previously, Tat transport has been investigated quantitatively in bacteria either using pulse-chase experiments *in vivo* (9) or using an inverted-membrane vesicle (IMV) transport system *in vitro* (10). Both methods are labor-intensive procedures since protein samples must be obtained over a time course followed by SDS-PAGE and autoradiography or immunoblotting. Additionally, such assays can only achieve a discontinuous monitoring of Tat transport over relatively large time intervals (mins); the acquisition of transport kinetics relying on fitting the time-course data with regression models. While such assays are sufficient for many issues, they are not well-suited for more advanced and comprehensive analyses, especially for monitoring rapid responses to different experimental conditions. Alternative procedures to investigate Tat transport have been proposed by quantifying the fluorescence of the green-fluorescent protein (GFP) bearing a Tat signal sequence. However, these studies were also performed with discontinuous measurements, GFP fluorescence either being quantified as end points using a photon-counting spectrophotometer (11, 12), or flow cytometry (13). Additionally, it is difficult to distinguish the fluorescence generated from the GFP precursor in the cytoplasm versus the mature GFP in the periplasm without further experimentation. Also, change of the cellular conformation such as formation of cell chains due to the defect in transporting the Tat substrates AmiA and AmiC (14) affects the quantitation of the GFP fluorescence measurement. Finally, transport of a completely alien substrate like GFP with a signal peptide from a Tat substrate may not reflect the Tat transport process using native cargo proteins. Thus, in order to address the limitations of the current transport assays and to conduct a more detailed analysis of Tat transport, it is important to develop a versatile, sensitive, and real-time Tat transport assay.

Recently, NanoLuc, a novel bioluminescence platform, has been developed and widely used for high-resolution imaging and other biological applications (15). Hallmarks of this system include its high stability, luminescence specificity and efficiency, and ATP-independence. Initially tailored for the study of protein-protein interaction, NanoLuc has been engineered to be split into two complementary components. The NanoLuc luciferase binary reporter system (NanoLuc Binary Technology, NanoBiT in short) (16) uses a small fragment, pep86 (1.3 kD, trademark name HiBiT), which interacts with the large component, 11S (18 kD, trademark name LgBiT), to generate a functional protein that luminesces with high intensity at 460 nm in the presence of its substrate furimazine (15). This NanoBiT system has a number of attractive features suggesting its utility for the development of a real-time Tat transport assay. First, previous studies have utilized the NanoBiT system to investigate protein secretion and transport in both mammalian cells and bacteria, both *in vivo* and *in vitro* (17–19). Second, in contrast to common split-luciferase systems which often require the presence of co-factors in addition to their substrate to produce luminescence, NanoBiT only requires molecular oxygen and its substrate furimazine to generate bioluminescence (15). This can minimize the potential interference of other reagents with the systems under investigation. Third, the rapid spontaneous interaction between the pep86-tagged Tat substrate and the 11S component (Kd of 0.7 x 10^-9^ M) generates luminescence allowing for the real-time monitoring of these two components in the same compartment (16). Finally, the enzymatic amplification of the luminescence signals by NanoLuc increases the sensitivity of the measurement (16), making it feasible to observe subtle changes upon different experimental treatments.

The Collinson group has exploited these properties utilizing the NanoBiT system to investigate protein transport via the Sec pathway in *E. coli* and protein import in mitochondria (17, 19, 20). In contrast to the Tat pathway, which transports folded proteins in an ATP-independent manner, the Sec and mitochondrial pathways transport proteins in their extended, unfolded conformations in an ATP-dependent manner (21). The NanoBiT system has allowed this group to measure elementary steps in bacterial protein export via the Sec pathway using proteoliposomes (PLs) incorporating the Sec machinery (17).

In this study, we have used NanoBiT technology to investigate transport via the Tat pathway. An *in vivo* NanoBiT-based real-time protein transport assay is reported using *E. coli* spheroplasts. Upon the validation of the Tat NanoLuc transport assay, we applied it to investigate the energetics of Tat transport by monitoring the real-time response of the transport signal to dissipation of different components of the PMF.

## Results

### Establishment of an *in vivo* real-time transport assay for the Tat pathway in *E. coli*

In order to develop a bioluminescence-based real-time assay using the NanoBiT system, we needed to transport a pep86-tagged Tat substrate into the compartment containing the 11S portion of the luciferase on the trans-side of the *E. coli* inner membrane (Figure 1A). Since the 11S polypeptide does not freely cross the bacterial outer membrane, we established the NanoLuc assay in spheroplasts, with the outer membrane having been removed. Transport in spheroplasts has been studied previously (22, 23), establishing some precedent for this approach. We attached the small fragment of the luciferase, pep86, to either the N- or C-termini of SufI (designated Npep86-SufI or SufI-Cpep86, respectively), a well-characterized bacterial Tat substrate (9, 24) (Figure 1B), and placed them under the control of T5-lacO promoter, which can be induced by the addition of isopropyl β-D-1-thiogalactopyranoside (IPTG). We then screened these pep86-tagged SufI constructs to determine their suitability for the transport assay. A conventional *in vivo* transport assay (25) was carried out to evaluate the performance of both pep86 variants by isolating the periplasmic fraction of cells co-expressing either wild-type TatABC (i.e., wtTat) or *ΔtatA,* together with SufI, SufI-Cpep86, or Npep86-SufI, followed by SDS-PAGE and immunoblotting (Figure 1C-E). Cells with the wild-type Tat expressing either SufI-Cpep86 or Npep86-SufI achieved more than 50% of transport relative to the wild-type Tat system transporting non-pep86-tagged SufI (Figure 1D). As a negative control, no detectable transport of SufI, SufI-Cpep86, nor Npep86-SufI was seen with the *ΔtatA* mutant, as TatA is an essential component of a functional Tat translocon. These results indicate that attachment of pep86 at either N- or C-terminal of SufI still allows for SufI to retain significant transport activity relative to native SufI. Next, we evaluated the level of overall protein induction by examining the total cell extract for each mutant. Cells expressing SufI-Cpep86 had a similar level of precursor production as the cells expressing native SufI. The expression level of Npep86-SufI, however, was significantly lower; approximately 1% relative to the level of SufI or SufI-Cpep86 (Figure 1E). Accordingly, SufI-Cpep86 was chosen as the most suitable substrate variant to investigate Tat transport via the NanoLuc system, as its transportability and overall expression level most closely resembles native SufI. Consequently, the following measurements were conducted using the cells expressing and transporting SufI-Cpep86.

**Figure 1.**
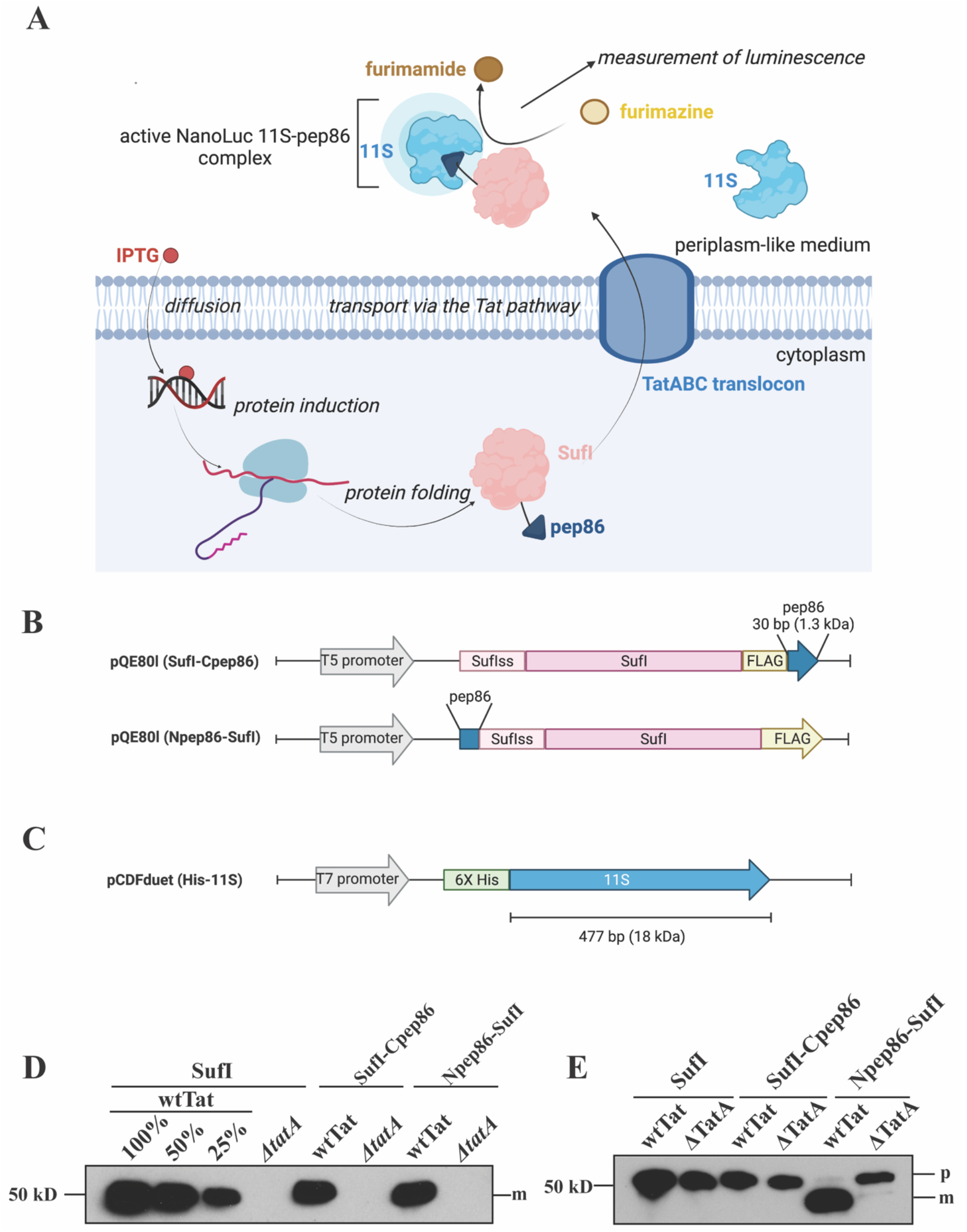
Schematic of *in vivo* Tat NanoLuc transport assay, and design of the pep86-variant selection. **(A)** Diagram explaining the experimental design of the NanoLuc transport assay. **(B)** Gene diagram for the pep86-tagged SufI variant. pep86 was added either at the N- (i.e., Npep86-SufI) or C- (i.e., SufI-Cpep86) terminus of the SufI. (**C**) Gene diagram for the 11S construct. A 6X His tag was added at the N-terminus for protein purification. (**D**) Assessment of the Tat transport activity among SufI-pep86 variants and native SufI. Periplasmic fractions from the *in vivo* transport assay were subjected to immunoblotting. (**E**) Assessment of the protein induction levels of the SufI-pep86 variants and native SufI. 1% of the total extract of the cells expressing native SufI or SufI-Cpep86 is shown in lanes 1-4, and 100% of the total extract of the cells expressing Npep86-SufI is shown in lanes 5 and 6. Position of the 50 kDa molecular weight marker is shown to the left; p, precursor; m, mature.

To set up the luminescence assay, we combined spheroplasts, overexpressed 11S, and the NanoLuc substrate furimazine in reaction medium. Upon the addition of IPTG, transcription and subsequent translation of the SufI-Cpep86 construct was induced. When this Tat substrate was transported out of the spheroplasts across the cytoplasmic membrane into the environment (i.e., 1X M9^+^ medium, which is the equivalent of periplasm), it came into contact with 11S, forming the active NanoLuc (11S-pep86 complex). Accompanied by the presence of furimazine and O2, it produced luminescence, which was monitored throughout the experimental time course (Figure 1A).

### The *in vivo* Tat NanoLuc Transport Assay faithfully reports Tat transport in spheroplasts

Next, we examined SufI-Cpep86 transport in spheroplasts of wtTat and the Tat-deficient mutant *ΔtatA* to confirm that the luminescence data correctly reflects Tat transport kinetics. Upon induction with IPTG followed by ~2 mins of lag time required for transcription and translation, a continuous increase in the luminescence signal for the wtTat spheroplasts was observed (Figure 2A). No significant increase in the luminescence signal was observed for the *ΔtatA* spheroplasts. To validate that the increase in luminescence corresponded to Tat transport, the periplasmic-mimetic external medium was cleared of spheroplasts by centrifugation and subsequently subjected to SDS-PAGE and immunoblotting. No mature SufI-Cpep86 was observed in the sample collected 1 min after IPTG induction. Mature SufI-Cpep86 was observed after 20 min in wtTat spheroplasts, whereas no mature protein was observed after 20 min induction in *ΔtatA* spheroplasts. Such results are consistent with the observed NanoLuc luminescence. Additionally, the amount of SufI-Cpep86 precursor induced was similar in the total cell extracts of the wtTat and *ΔtatA* spheroplasts, which indicates that the difference in the luminescence signal between the wtTat and *ΔtatA* spheroplasts was due to a difference in transport rather than the lack of SufI-Cpep86 production in the absence of TatA (Figure 2A). Notably, a very small fraction of precursor was present in the periplasmic samples from the *ΔtatA* spheroplasts. Previous studies have reported that, due to the absence of the Tat activity, *ΔtatA* spheroplasts are more susceptible to lysis compared to the wtTat cells (26). Nevertheless, the amount of precursor present in the external medium is quite small and negligible relative to the total amount of precursor synthesized (Figure 2A).

**Figure 2.**
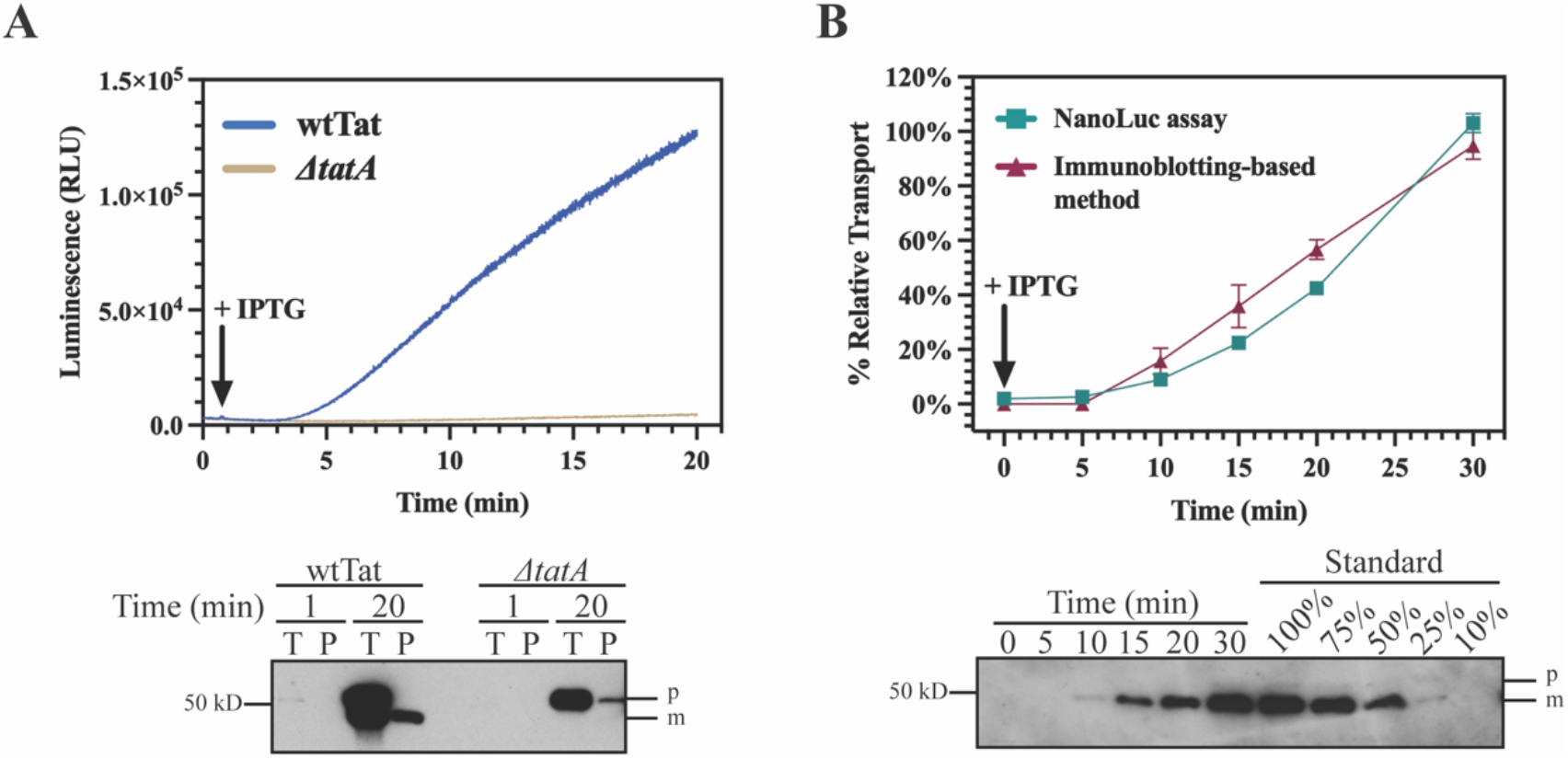
The NanoLuc assay gives a genuine measure of Tat transport. (**A**) The NanoLuc assay correctly reports Tat-specific transport. wtTat transporting SufI-Cpep86 as the positive control (red curve); *ΔtatA* is shown as the negative control (blue curve). Arrow indicates the time when IPTG was added protein induction. Lower panel: samples were taken from the reaction cuvette after either 1 or 20 min from the reaction cuvette and immunoblotted using an antiFLAG-tag antibody. (**B**) The NanoLuc assay yields results consistent with the conventional immunoblotting-based method. Transport is normalized based on the level of transport at 30 min (100%). Error bars indicate standard error from three replicates; representative immunoblot shown in the lower panel. RLU, relative light units; p, precursor; m, mature. Other details as in Fig. 1.

In addition to confirming that the NanoLuc luminescence measurement records the SufI transport kinetics in a Tat-specific manner, we evaluated the consistency between the kinetic curves obtained from the luminescence assay and those from the conventional immunoblotting-based method. Three technical replicates were performed to assess the reproducibility of the NanoLuc assay. Upon the IPTG induction, samples were removed at six different time points, 0, 5, 10, 15, 20, and 30 min post-IPTG induction. For the luminescence assay measurement, the corresponding luminescence readings were recorded from these samples (Figure 2B). For the conventional method, samples were collected at the same time points and placed on ice to stop transport. These samples were then subjected to SDS-PAGE followed by immunoblotting, and the transport kinetics were analyzed (Figure 2B). These results demonstrated that the kinetic curves obtained from the luminescence assay fully recapitulated those obtained from the conventional immunoblotting-based method. In addition to this consistency, the NanoLuc assay exhibits the following merits. First, the variation among replicates in NanoLuc assay is relatively small compared to the conventional method, yielding measurements with higher resolution. In this case, the luminescence assay allows us to identify small but significant differences in transport between different experimental treatments. Second, the luminescence measurement in the NanoLuc assay is in real time, which can minimize the experimental errors during sample handling. For example, it is observed that the percentages of transport relative to the end point were higher using the conventional method relative to the luminescence assay. Such overestimation of the transport was perhaps due to inefficient quenching of the transport process by ice, a problem rendered moot in the real-time NanoLuc assay.

Taken together, our results indicate that the NanoLuc assay is a legitimate measure of Tat transport activity that is continuous, exhibits high resolution and reproducibility, and which is therefore a suitable method for comprehensively investigating the Tat transport kinetics described in the forthcoming section and in future studies.

### Investigation of the protonmotive force dependency of the Tat machinery in *E. coli* spheroplasts

The Tat transport pathway was originally discovered in thylakoids in part based on its requirement for a ΔpH and lack of requirement for ATP to power protein transport (27). Whilst valinomycin was initially observed to be without effect on the protein transport, an effect of the Δψ was ultimately observed under limiting light intensity, indicating that in thylakoids the Tat pathway was powered by both components of the PMF (6). Bageshwar and Musser studied the energy requirement for Tat transport into inverted membrane vesicles derived from *E. coli* and concluded that it was driven by the Δψ only, with no contribution from a ΔpH (10). We thought it was unlikely that two different mechanisms governed the same transport process in different organisms and recognized that our newly developed real-time luminescence assay afforded us the opportunity to revisit this question.

With these considerations in mind, we applied the NanoLuc assay to investigate Tat transport energetics in spheroplasts with respect to the different components of the PMF. First, we examined the response to collapse of the Δψ by the addition of 1 – 2 μM valinomycin (V) (Figure 3A). As expected, the transport-reporting luminescence curves plateaued after valinomycin addition, while the positive control in which only the solvent ethanol was added (E) continued to increase unabated. This is consistent with the known requirement for the Δψ for Tat transport in *E. coli*. We next observed that addition of 1 – 10 μM nigericin also caused the rising luminescent signal to plateau, suggesting that, as in thylakoids, the ΔpH also was able to drive protein transport. This finding differed from the conclusions reached earlier in which nigericin was not observed to inhibit Tat transport in IMVs (10).

**Figure 3.**
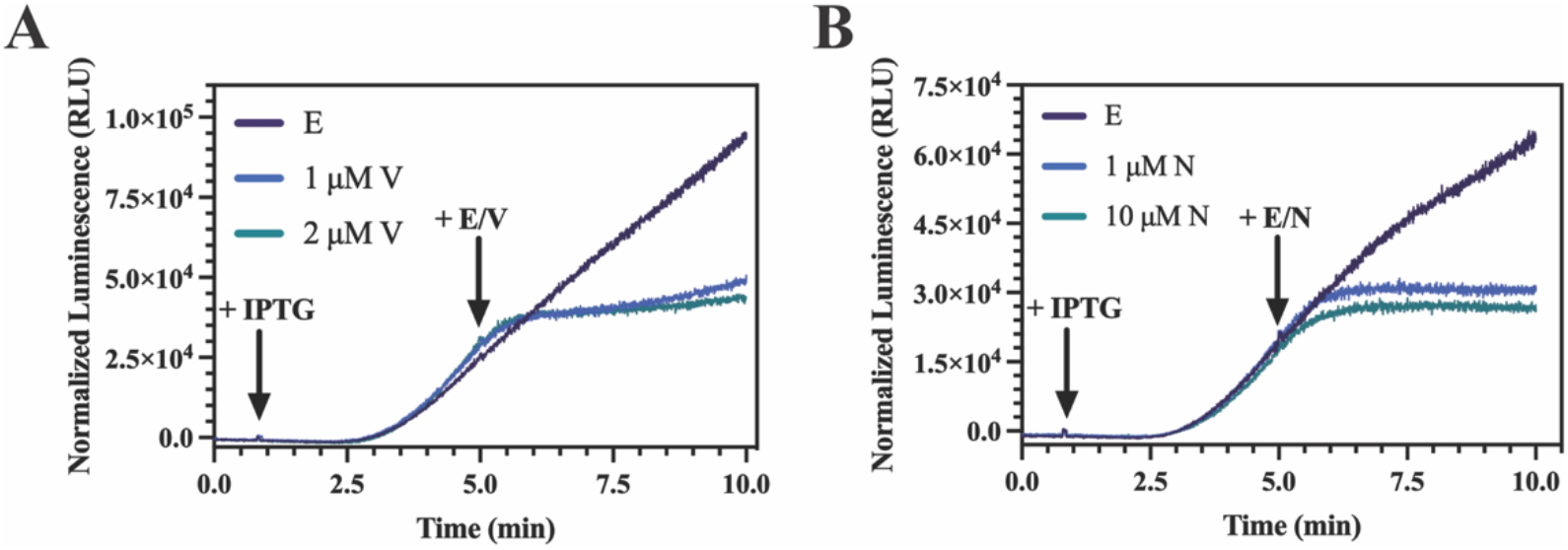
Both ΔpH and Δψ appear to play a role in Tat transport in *E. coli* spheroplasts. Transport curves with SufI-Cpep86 in *E. coli* spheroplasts are shown. A background signal was calculated by averaging the luminescence before IPTG addition, and this background is subtracted from the luminescence signals. IPTG was added at the first arrow, ionophores (valinomycin (V), or nigericin (N)) or their solvent (ethanol (E)) were added at the second arrow. Other details as in Fig. 2.

To further investigate our discrepant result, we examined several artefacts that could potentially result in the loss of the increasing NanoLuc signal upon addition of nigericin or valinomycin. First, it is possible that nigericin and/or valinomycin, themselves, could negatively impact the 11S-pep86 interaction and, consequently, affect the NanoLuc signal. To assess this possibility, we examined the direct effect of nigericin and valinomycin on the 11S-SufI-Cpep86 interaction. To this end, excess purified 11S and furimazine were placed in 1X M9^+^ medium (without *E. coli* spheroplasts) and SufI-Cpep86 was added at t = 20 sec to initiate the NanoLuc luminescence reaction. When the luminescence signal reached the maximum (~160 sec), ethanol, valinomycin, or nigericin were added to the indicated final concentrations either individually or together to assess their effects on the NanoLuc signal. As can be seen in Fig. 4, these additions caused a dose-dependent decrease in the NanoLuc luminescence signal, indicating some detrimental interaction between the ionophores and the luciferase. At 220 sec, same amounts of purified SufI-Cpep86 was again added to determine whether the presence of the ionophores irreversibly hindered the interaction between 11S and SufI-Cpep86; it did not. Notably, ethanol alone caused a reproducible decrease in the NanoLuc signal (Figure 4A-C). Accordingly, we expressed in Fig. 4D the inhibition of the NanoLuc signal as a percentage of the nigericin or valinomycin treatment relative to the ethanol treatment to remove the background effect of the solvent alone. A cutoff of 90% was used to assess whether the ionophore effect is significant (Figure 4D). Valinomycin at a final concentration of 1 μM or higher exhibited a significantly negative impact on the NanoLuc signal, such that 1 μM brought the signal to ~68%, and 2 μM lowered it to ~48% relative to the positive ethanol control (Figure 4A and 4D). For nigericin, 10 μM dropped the signal to ~43% of the ethanol treatment. Interestingly, 1 μM of nigericin appeared to minimally mitigate the effect of ethanol on the NanoLuc signal, such that it produced a value higher than 100% relative to the ethanol treatment (Figure 4B and 4D). While we do not currently know why these ionophores negatively affected the NanoLuc signal and did not pursue it further, we noted that their effects were minimal when their concentrations remained below 0.5 μM, separately or in combination, a concentration still expected to dissipate the PMF in spheroplasts (Figure 6).

**Figure 4.**
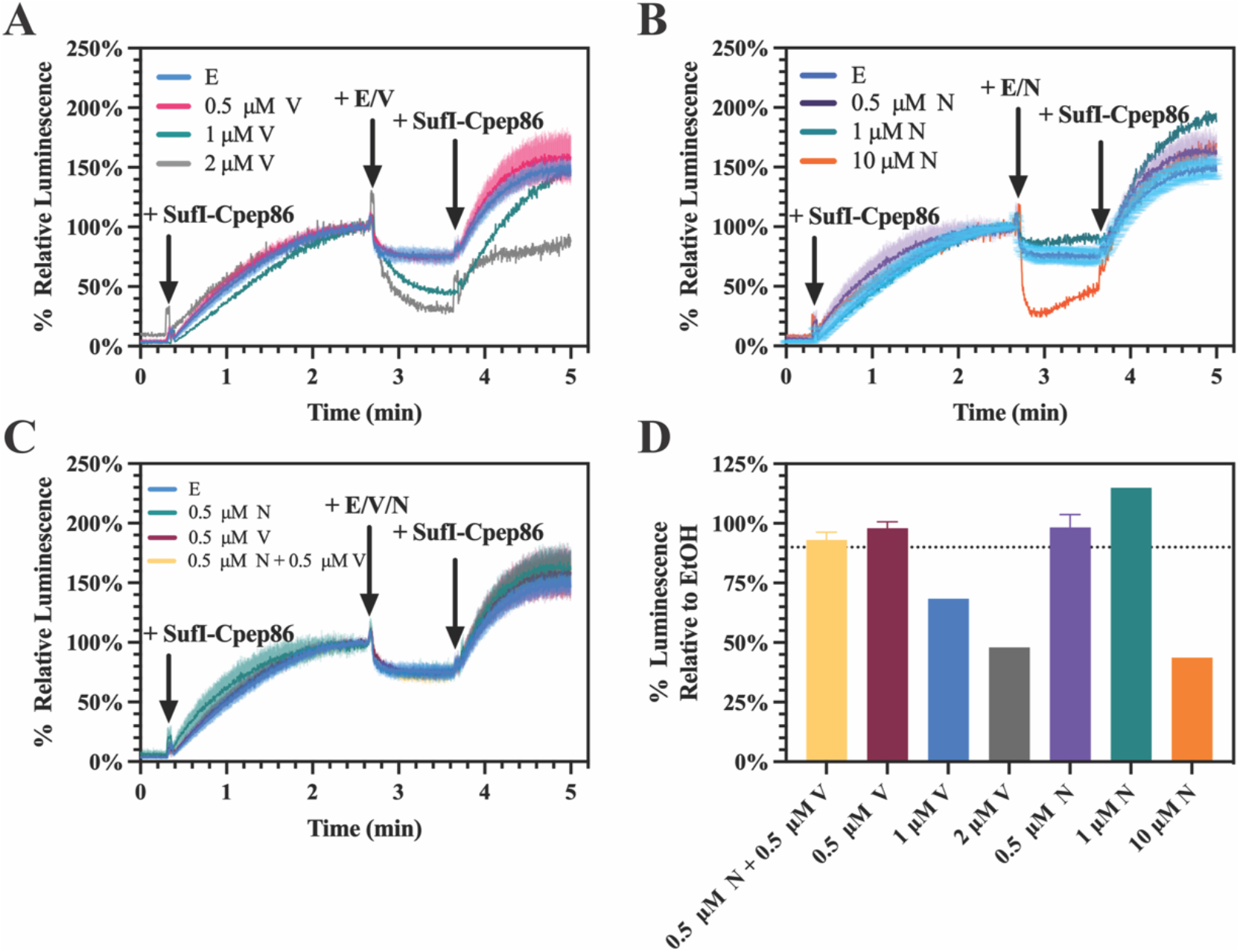
Effects of the ionophores on the NanoLuc signals. (**A-C**) NanoLuc luminescence in the presence of ethanol (E), valinomycin (V), nigericin (N), or both ionophores (N+V) at the indicated final concentrations. Purified SufI-Cpep86 was added to a final concentration of 6.6 nM. (**D**) Bar plot showing the percent luminescence upon ionophores addition relative to the ethanol addition. A cutoff of 90% is denoted as dotted line. Error bars represent standard error among three technical replicates.

In addition to nigericin or valinomycin directly affecting the NanoLuc signal, it is possible that they could also exhibit a negative impact on the synthesis of SufI-Cpep86 precursor, and this could also prevent the luminescence signal from increasing in the NanoLuc assay. Previous studies have shown that ionophores, including valinomycin and nigericin, mildly inhibited protein synthesis in *E. coli* but exhibited a stronger protein synthesis inhibition in the gram-positive bacteria *S. aureus* which lacks outer membranes (28). Other studies have indicated that the *E. coli* outer membrane serves as a barrier against the entry of ionophores into the cell (29). Accordingly, we could not rule out the possibility that nigericin and valinomycin could inhibit protein synthesis in the *E. coli* spheroplasts as these are devoid of the outer membrane. In order to test this possibility, a spheroplast Tat transport assay was carried out in which we monitored the presence of both precursor and mature substrate (Figure 5). IPTG was added to the wtTat spheroplasts to induce the expression of SufI-Cpep86 at 0 min, followed by the addition of ethanol (E), nigericin (N), or valinomycin (V), either individually or together. Transport reactions continued for the next 15 min before harvesting the spheroplasts. Both precursor and mature SufI-Cpep86 were observed in the ethanol treatment after 15 min, indicating successful protein induction and transport. However, neither mature nor precursor SufI-Cpep86 were observed after addition of nigericin or nigericin plus valinomycin, which suggests that nigericin strongly impedes the induction of protein synthesis. It should be noted that, although faint precursor bands were observed in the valinomycin treatment, these were not comparable to the ethanol treatment (Figure 5A). Taken together, these results indicate that nigericin and valinomycin at concentrations above 1 μM negatively affect the synthesis of the SufI-Cpep86 protein precursor in *E. coli* spheroplasts. In this case, we could not distinguish whether a flattening in the luminescence signal in the NanoLuc assay upon ionophore addition (Figure 3) was due to cessation of transport or to an inability of the cells to replenish the precursor protein.

**Figure 5.**
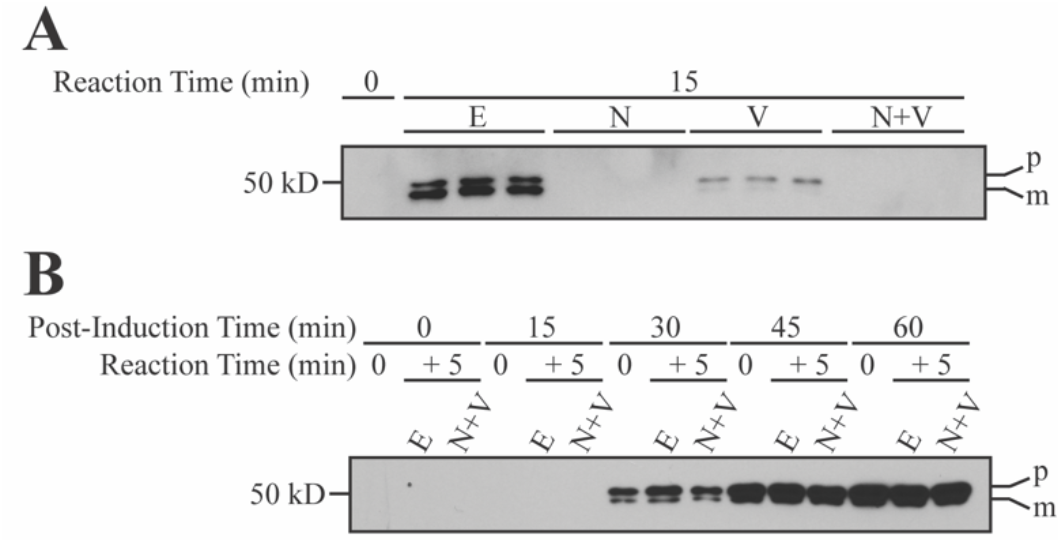
Effects of ionophores on protein synthesis. Total extracts of *E. coli* spheroplasts expressing wtTat and FLAG-tagged SufI-Cpep86 were immunoblotted. (**A**) Ethanol or ionophores were added at 0 min and reactions continued for 15 min. Ethanol (E); 0.5 μM valinomycin (V); 0.5 μM nigericin (N); 0.5 μM of both ionophores (N+V). (**B**) Top row indicates the amount of time in minutes after IPTG induction. Second row indicates the time before ethanol or ionophores were added (i.e., 0 min) and after the reaction (i.e., 5 min). A representative immunoblot from three replicates is shown. Other details as in Fig. 2.

In order to minimize the negative impact on protein synthesis upon ionophore addition, we needed to identify a time window for the experiment which satisfies the following criteria. First, sufficient amounts of SufI-Cpep86 precursor should be present throughout the measuring time window so that refilling of the precursor pool by new synthesis is not required. Second, the measuring time should be short enough that minimal changes occur in the total amount of precursor during the reaction. Upon fulfilling these conditions, the possibility that a lack of precursor leads to a flattening of the NanoLuc curve can be ruled out. To determine the optimal time window, IPTG was added to the wtTat spheroplast to initiate precursor synthesis at 0 min, followed by the addition of ethanol (E) or nigericin plus valinomycin (N+V) at t = 0, 15, 30, 45, and 60 min post-IPTG induction. The transport reactions were then continued for the next 5 min, after which the spheroplasts were harvested. We then compared the level of protein precursor before and after the various treatments. As seen in Fig. 5B, detectable amounts of SufI-Cpep86 precursor were observed 30 min after IPTG induction. However, the overall amount of precursor present was rather low, and it decreased in the 5 min after ionophore addition (compare lanes E and N+V at 30 min). However, after 45-min post induction, there is minimal visual difference in the level of precursor protein between the E and N+V samples, and before the additions (Figure 5B). Based on these results, we chose to monitor the transport reactions for 5 min at 45 min post-induction as the optimal time for ionophore addition to ensure the presence of a sufficient amount of SufI-Cpep86 precursor.

Consequently, we modified the NanoLuc assay to incorporate these optimized conditions. To wit, the spheroplasts were incubated for 45 min after IPTG induction to allow the accumulation of an excess of precursor. Subsequently, spheroplasts were pelleted and resuspended to remove the accumulated mature SufI-Cpep86 transported out of the cells to the medium, which reduced the background signal. Washed spheroplasts were resuspended with fresh 1X M9^+^ medium, and then the NanoLuc assay was initiated. First, we performed the NanoLuc assay without further adjusting the pH of the medium (pHout = 6.3). Ethanol (E), nigericin (N), valinomycin (V), or both ionophores (N+V) were added at 20 sec. The curve for ethanol treatment continued to increase over the 5 min monitoring time, indicating continuous protein transport. Upon addition of both ionophores or valinomycin alone, the signal reached a plateau around 3.5 min, i.e., the transport reaction stopped, consistent with the requirement for the Δψ for protein transport observed earlier (10). Notably, addition of nigericin alone also caused the transport reaction to stop, with the NanoLuc signal behaving as was observed when valinomycin was added; that is, transport stopped by approximately 3.5 mins (Figure 6A). Given that nigericin at concentrations as low as 0.05 μM have been observed to be sufficient to abolish the ΔpH in the outer-membrane deficient *E. coli* cells (29), we expect that the 0.5 μM used in this experiment effectively dissipated the ΔpH in our spheroplasts. Considering the possibility that the change in the NanoLuc signals is an artefact of a significant change in the amount of spheroplasts present throughout the measurement, we compared the spheroplast-to-whole-cell ratio before and after the measurement using the spheroplast lysis method described in Materials and Methods (23). Regardless of the addition of ethanol or ionophores, the percentages of spheroplasts after the measurement averaged greater than 90% of the spheroplast percentages before the measurement (Supplemental Figure 2). Hence, it is unlikely that the significant changes observed upon the addition of ionophores are due to the changes in the number of spheroplasts present in the reaction. Accordingly, the separate inhibition of Tat protein transport in *E. coli* by both nigericin and valinomycin indicates that this reaction is powered by both components of the PMF, as is the case for thylakoids (6).

**Figure 6.**
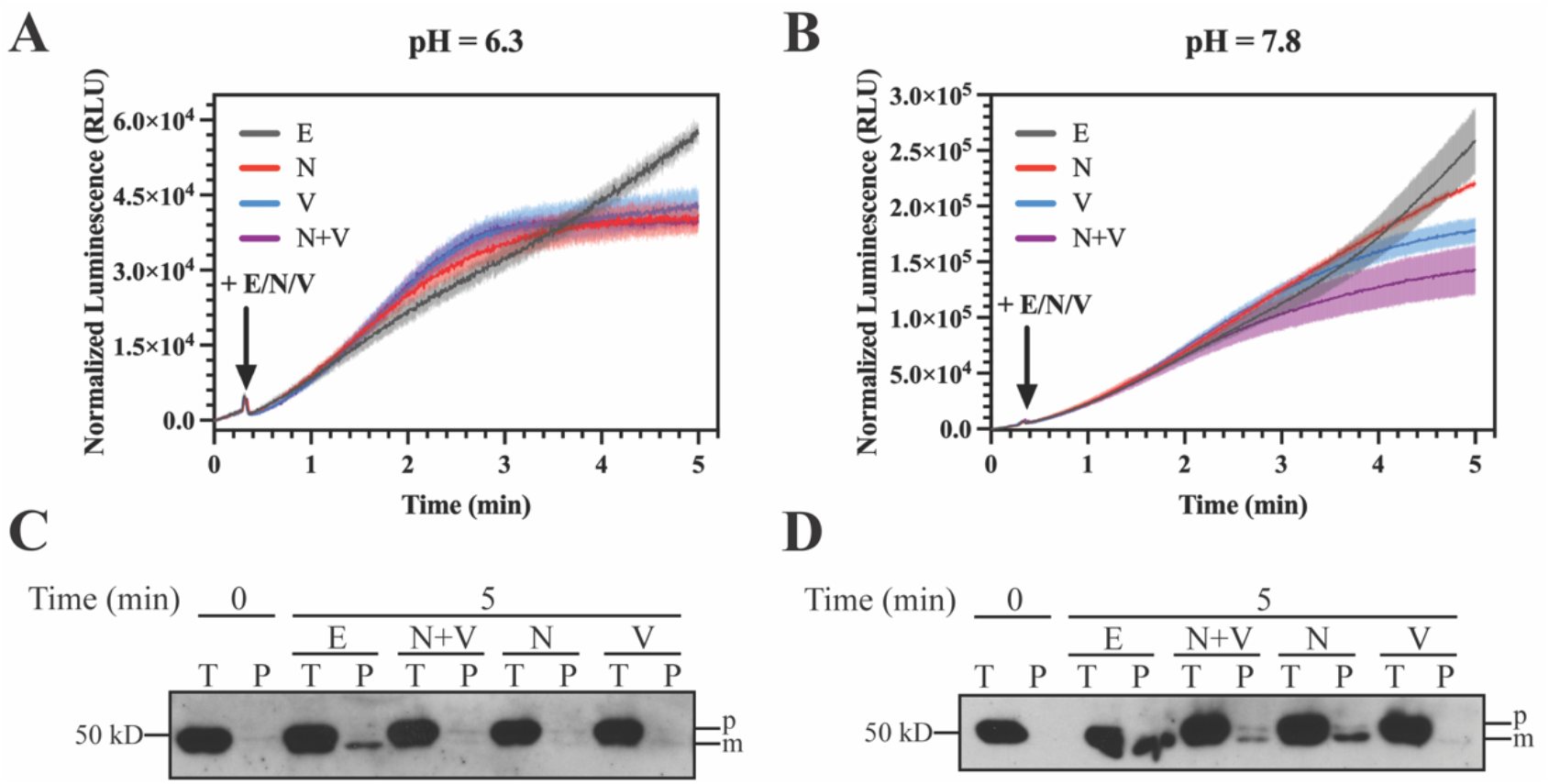
The NanoLuc assay reports a role for ΔpH in Tat transport in *E. coli* spheroplasts. (**A-B**) NanoLuc transport curves at pH = 6.3 (**A**) and pH = 7.8 (**B**). Arrows indicate addition of ethanol or ionophores. Shades represent the standard error from three replicates. E, ethanol; N, 0.5 μM of nigericin; V, 0.5 μM of valinomycin; N+V, 0.5 μM each nigericin and valinomycin. (**C-D**) Representative immunoblots of the samples taken at the beginning (0 min) and the end (5 min) of the NanoLuc assay at pH = 6.3 (**C**) and pH = 7.8 (**D**). T, total extract; P, periplasmic fraction (i.e., supernatant); other details as in Fig. 2. Three replicates were performed for each group. Other details as in Fig. 2.

To further examine the effects of nigericin on Tat transport in our experiments, we carried out the same NanoLuc assay at an elevated pH of 7.8 in the external medium. i.e., closer to that of the cytoplasm (30). We reasoned that at this higher pH, the spheroplasts would be unable to develop a significant ΔpH between the cytoplasm and the periplasmic-like external medium. Under these conditions we might expect the effect of nigericin to be minimal if its principal action in earlier experiments was to dissipate a ΔpH. Figure 6B shows that this was indeed the case. As in Figure 6A, at pH 7.8 (Figure 6B) valinomycin, either alone or combined with nigericin, continued to cause cessation of the rise of the NanoLuc signal. However, the effect of nigericin addition was greatly diminished at this pH. It is noteworthy that the luminescence produced by the recombined luciferase is significantly higher at pH 7.8 than at pH 6.3, which we expected due to our observed enhanced NanoLuc activity at higher pH (Supplemental Figure 1). Nonetheless, the relative slopes of the control curves were similar. We can conclude from this experiment that the effect of nigericin observed at pH 6.3 was due to its dissipation of the ΔpH and not an effect on the luciferase reaction.

We also examined by SDS-PAGE and immunoblotting the total cell extract and the periplasmic samples which were collected before and after these luminescence measurements. At pH 6.3, only the ethanol treatment exhibited a strong mature band (Figure 6C), and valinomycin and nigericin both inhibited protein transport. At pH 7.8, nigericin caused much less inhibition of Tat transport (Figure 6D), consistent with the idea that the ΔpH contributes only minimally to the PMF at this high pH. Additionally, the level of precursor present in the different treatments at the end of luminescence measurements were similar, which was consistent with our previous results. Notably, we observed faint background bands of the precursor in the periplasmic samples. This may have been caused by a small amount of lysis of the spheroplasts during the measurement. However, the level of the background was much lower than the ethanol treatment, which was the positive control (Figure 5B, and Figure 6 C-D).

Taken together, we have modified the NanoLuc assay to report the transport kinetics via the Tat pathway in *E. coli* spheroplasts. Consistent with previous findings *in vitro* using *E. coli* IMVs (10), dissipation of the bulk PMF by adding nigericin and valinomycin, together, as well as dissipation of the Δψ alone, using valinomycin, abolishes Tat transport. However, we observed that ΔpH also plays a role in the Tat transport in *E. coli* spheroplasts *in vivo* using the NanoLuc assay. Tat transport was stopped when the proton gradient was dissipated by nigericin under the conditions where a significant ΔpH (pH_out_ = 6.3) was present. It follows then that the energetics of the Tat pathway in bacteria is not substantially different from that in thylakoids, relying on both components of the PMF.

## Discussion

In order to achieve a better understanding of the Tat transport mechanism, a continuous assay to monitor the transport with high resolution is desirable. Lack of such an assay makes it difficult to investigate Tat transport kinetics in a comprehensive manner. Current quantitative assays in investigation of the Tat transport *in vivo,* such as pulse-chase experiments, requires a series of complex procedures and relies on fitting with a regression model to estimate the transport kinetics (9). Here, we describe and validate a novel luminescence Tat transport assay utilizing the split-NanoLuc luciferase system to monitor the Tat transport kinetics *in vivo.* Merits of the NanoLuc assay include the following considerations: First, by using *E. coli* spheroplasts as the experimental platform, the Tat transport is assessed *in vivo*, under nearly native conditions. Second, since the Tat machinery transports proteins in their folded conformations, disruption of the original conformation upon tag addition to the Tat substrate should be minimized or avoided. This is achieved with the NanoLuc complement pep86 (10 aa), where its impact on the overall conformation of the Tat substrate is expected to be relatively small. Third, the sensitive and continuous nature of the NanoLuc assay allows us to observe the transient and subtle changes in transport upon the application of treatments during the reaction time course. Fourth, the NanoLuc assay involves relatively few steps compared to conventional pulse-chase experiments and *in vivo* transport assays, minimizing sample handling. While our results and previous studies (16) indicate that pH and several ionophores have a direct effect on the NanoLuc signals (Figure 4 and Supplemental Figure 1), we were able to manage these effects by optimization of the assay. Altogether, our results indicate that the NanoLuc assay is a convenient and accurate measure of Tat transport activity.

The energy requirement of the Tat transport system in both bacteria and chloroplasts has been studied for decades (6, 10, 27). In plants, ΔpH is the dominant driving force for Tat transport across the thylakoid membrane (27, 31, 32). This is likely because thylakoids generally partition most of their PMF as a ΔpH, although a substantial with a substantial Δψ can be developed under some conditions, both *in vivo* and *in vitro* (33). Indeed, thylakoids can utilize the Δψ to drive Tat protein transport, and accordingly, the thylakoid PMF is considered to power this process (6). In contrast to this, it has been suggested that the Δψ, but not ΔpH, drives Tat protein transport in *E. coli* (10). This was surprising given that the Tat pathway has long been thought to operate by a conserved mechanism in both plants and bacteria (1).

We recognized that our new luminescence assay for Tat transport in *E. coli* affords us a new opportunity to examine the energetics of this process in bacteria *in vivo.* Under our conditions and using *E. coli* spheroplasts, we report that the ΔpH, in addition to the Δψ, plays a role in the Tat transport. When a significant ΔpH was allowed to develop by electron transport, dissipation of the proton gradient using nigericin resulted in a rapid abolishment of Tat transport. Similarly, and as has been reported earlier with in vitro studies (10), addition of valinomycin also lead to inhibition of Tat transport, confirming the role of the Δψ as a driving force. (Figure 6A and 6C). Our results, which indicate that both ΔpH and Δψ play a role in the Tat transport, unifies the PMF-dependency of the Tat transport in both thylakoids and bacteria. It should be noted that previous studies used *E. coli* inverted membrane vesicles (IMVs) as the experimental platform (10). This may be due in part to the size difference between the membrane vesicles (~100 nm in diameter) and *E. coli* spheroplasts (~ 1 μm in diameter), resulting in a much smaller periplasmic volume in IMVs (34, 35).

In our experience, and as was already noted in the study of Bageshwar and Musser, we find it difficult to assess the energetic parameters operating during the entirety of the Tat transport reaction in IMVs. Addition of NADH to IMVs permits a measurement of the developed △pH and △ψ for only a few minutes until the reaction vessel is depleted of oxygen. However, the Tat transport reaction continues after components of the PMF can no longer be detected (10). While we cannot measure △pH or △ψ in spheroplasts, we do recognize that the energetic profile in living cells could be distinctly different than in IMVs. In particular, even if the spheroplast reaction vessel were to go anaerobic during the experiment, we could expect the PMF driving force for Tat transport to be maintained by reverse proton pumping by the EF1/EF0 ATPase using glycolysis-derived ATP. Furthermore, taking the possible compensation in Δψ upon nigericin addition into account, we have optimized the concentration of nigericin used in this study (0.5 μM), which is within the range (0.05 – 2 μM) found to be sufficient to dissipate the ΔpH in *E. coli* but to leave the Δψ remain relatively unchanged (29). Still, we observed that Tat transport stopped when ΔpH was dissipated by nigericin at pH = 6.3, suggesting a role of ΔpH in the Tat transport (Figure 6A and 6C). To further support these observations, we performed the same set of experiment at the elevated pH of 7.8, where the environmental pH is close to the *E. coli* cytoplasmic pH (30). We found that addition of nigericin had minimal effect on the Tat transport under these conditions (Figure 6B and 6D), ruling out the possibility that Tat transport was stopped by nigericin at pH 6.3 due to some non-specific effect of the protonophore. We are left with the conclusion that the ΔpH contributes to the driving force for Tat transport in *E. coli* in vivo.

To summarize, we have examined bacterial Tat transport energetics *in vivo* using our novel high-resolution real-time assay with the NanoLuc split-luciferase system. Our experiments clearly demonstrate that both Δψ and ΔpH can contribute to the driving force for Tat transport, suggesting that this reaction is powered by the PMF. Accordingly, we propose that there is no need to consider that this enigmatic transporter operates by different principles or different mechanisms in the several domains of life where it is found.

## Materials and Methods

### Strain and Plasmid Construction

*E. coli* strain DADE-A (MC4100, *ΔtatABCE)* was used in in both *in vivo* transport assay and the luminescence assay (36). The plasmid pTat101 (pTH19Kr derivative, a low copy plasmid constitutively expressing the TatABC) (37) was used for Tat expression. *ΔtatA,* the TatA knockout variant, was produced by QuikChange site-directed mutagenesis (NEB, Phusion High-Fidelity PCR Kit) followed by KLD (Kinase, Ligase, and DpnI) digestion (NEB) by using primers deltaTatA_F (5’-GTGTTTGATATCGGTTTTAGC-3’) and deltaTatA_R (5’-GGATCCTCCTCTGTGGTA-3’). pep86 was introduced at the relevant position of the pQE80l (SufI-FLAG) by QuikChange site directed mutagenesis followed using the following primers: Npep86_F (5’-CCTGTTTAAAAAAATTGGCTCCGGCTCACTCAGTCGGCGT-3’) and Npep86_R (5’-CGCCAGCCGCTCACCATGGTTAATTTCTCCTCTTTAATGAATTCTG-3’) for N-terminal addition; Cpep86_F (5’-GCGCCTGTTTAAAAAAATTTAAGTCGACCTGCAGCCA-3’) and Cpep86_R (5’-CAGCCGCTCACGCCGGAGCCCTTATCGTCGTCATCCTTGTAATC-3’) for C-terminal addition. For the pCDFduet (His-11S) construct, 11S was synthesized with Ndel and Xhol restriction enzyme recognition sites and inserted into an empty pCDFduet under the control of the T7 promoter. 6X His tag was introduced by QuikChange mutagenesis using Nhis11S_F (5’-CATGGCGTCTTCACCCTGGAAGAC-3’) and Nhis11S_R (5’-GTGATGGTGACCCATATGTATATCT-3’). All constructs were confirmed by Sanger sequencing.

### *In vivo* Transport Assay

Plasmids pTat101 or *ΔtatA* were co-transformed with pQE80l (SufI-FLAG), pQE80l (Npep86-SufI-FLAG), or pQE80l (SufI-Cpep86-FLAG), which is under the control of the T5 promoter, into the Tat knockout strain, DADE-A. Details of the *in vivo* transport assay was described elsewhere (25). Overnight cultures were diluted into the fresh LB medium. 1 mM IPTG was added for induction of SufI and SufI-pep86 variants when the OD_600_ of the cell culture reached 0.5. Cells were continuously cultured at 37 °C for another 2.5 hrs before proceeding to cellular fractionation.

### Protein Expression and Purification

6X His-tagged 11S was cloned into the expression vector pCDFduet and transformed into BL21 (λDE3). When OD_600_ reached 0.7, proteins were overexpressed overnight at 4 °C upon induction with IPTG induction at a final concentration of 0.25 mM. Cells were placed on ice for 10 min before being centrifuged at 10,000 g for 10 min at 4 °C. Proteins were purified under native conditions by Ni-NTA chromatography in the following steps (10). Pellets were resuspended on ice in lysis buffer (25 mM CAPS, pH = 9, 250 mM NaCl, 20 mM imidazole, and 0.2% Triton X-100) to obtain a 50X concentrated cell lysate. Lysates were sonicated followed by centrifugation at 10,000 rpm (Beckman JA-14) for 25 min at 4 °C. Subsequently, for 10 mL cell lysates, 2.5 mL of the 50% Ni-NTA (Qiagen) slurry was added and gently mixed on a rotary shaker at 4 °C for 1 hr. The resin was then loaded onto a 10 X 3 cm column. The following buffers were added sequentially: (1) 15 mL wash buffer 1 (100 mM Tricine, pH = 8, 1 M NaCl, 20 mM imidazole, and 0.2% Triton X-100); (2) 15 mL wash buffer 2 (10 mM Tricine, pH = 8, 100 mM NaCl, and 10 mM imidazole); (3) 5 mL wash buffer 3 (10 mM Tricine, pH = 8, 100 mM NaCl, 10 mM imidazole, and 50% glycerol). 6X His-tagged proteins were eluted by adding 12 mL of elution buffer (100 mM NaCl, 250 mM imidazole, pH = 8.0, and 50% glycerol) and collected as 2-mL fractions. Buffer exchange was performed to exchange the elution buffer with protein storage buffer (50 mM Tris-HCl, pH = 8, 100 mM KCl, and 10% glycerol) using Amicon™ centrifugal filter units (MilliporeSigma) for long-term storage at −80 °C. Protein concentrations were estimated by measuring the absorbance at 280 nm and by SDS-PAGE.

### Preparation of the *E. coli* Spheroplast

Plasmids pTat101 or *ΔtatA* were co-transformed with pQE80l (SufI-Cpep86-FLAG) into the DADE-A strain. Overnight bacterial cultures were grown in LB medium supplemented with 0.4% glucose to suppress the expression of SufI-Cpep86. The next day, these cultures were diluted in fresh LB medium supplemented with 0.4% glucose at a ratio of 1:33 and grown at 37 °C until the OD_600_ reached 0.6-0.9. Preparation of spheroplast was derived from the protocol from Schäfer et al.(38). All centrifugations were performed at 5,000 rpm (Beckman JA-14) for 10 min at 4 °C. 200 mL of the cells with OD_600_ equivalent to 1.0 were centrifuged and resuspended with 45 mL of resuspension buffer (100 mM Tris-HCl, pH = 7.5, 0.5 M sucrose). Cells were converted to spheroplast by adding an equal volume of spheroplast-converting buffer (8 mM EDTA, pH = 8.0, 0.01 mg/mL lysozyme) and incubation on ice for 10 min. Assessment of the conversion percentage was carried out by calculating the ratio of OD_600_ of the samples diluted with ddH2O to the ones diluted by osmotic-stabilizing 1X M9^+^ medium (1X M9 medium diluted from 5X M9 salt, supplemented with 0.1 mM CaCl2, 0.002% thiamine, 2 mM MgSO4, 0.01% of the 20 amino acids, 2% glycerol as the carbon source, and 0.25 M sucrose) both at the ratio of 1:10. Since the spheroplasts lyse in ddH2O but not in 1X M9^+^ medium, this calculation yields the fraction of the cells remaining unconverted (23). The typical conversion rate from whole cell to spheroplast using this method was above 90%. The spheroplasts were centrifuged and washed with 1X M9^+^ medium once to remove the residual EDTA and lysozyme. Washed pellets were resuspended with 5 mL of 1X M9^+^ medium and the OD_600_ of the spheroplast mixture was measured. The concentrated spheroplast solution was incubated on ice and ready for use.

### Tat NanoLuc Transport Assay

The real-time standard luminescence assays were carried out at 37 °C in a Jobin Yvon Fluorolog (Horiba) fluorimeter with the excitation lamp turned off. The luminescence was measured with the emission wavelength set at 460 nm with the slits open with a 10 nm bandpass. The reaction mixture, with the total volume of 2 mL, was assembled in a 3.5 mL standard cuvette with a stir bar and contained the following components: concentrated spheroplast solution in 1X M9^+^ medium added to the final OD_600_ of 1.0, 50 nM His-11S, 2.5 μL furimazine (Promega), with the addition of 1X M9^+^ medium to a 2 mL volume. After equilibrating for 5 min, the luminescence baseline was measured. IPTG was then added after 50 sec to a final concentration of 1 mM. After a measurement for either 400 sec or 600 sec, the ionophore dissolved in ethanol or ethanol, alone, was added to the reaction mixture as indicated, and luminescence was continuously measured until completion of the reaction.

For the experiments in investigation of the Tat energetics, *E. coli* spheroplasts were diluted in 1X M9^+^ medium to the final OD_600_ of 0.5. The spheroplast premix was incubated at 37 °C with continuous stirring for 5 mins. IPTG was added to the final concentration of 1 mM. Spheroplasts were then incubated for 45 mins with stirring. The transport reaction was then paused by incubation on ice. Spheroplasts were washed by centrifugation at 5,000 rpm (Beckman JA-14) for 10 min at 4 °C to remove the excess transported SufI-Cpep86 in the periplasmic-like medium. Fresh 1X M9^+^ medium was used to resuspend the spheroplasts to the final OD_600_ of 0.5. The reaction mixture was prepared in the same manner as described above. Without further incubation, ethanol or ionophores were added after 20 sec. Luminescence was then measured continuously for 5 mins.

### Cellular Fractionation

For samples from the *in vivo* transport assay, 3 mL of cells with the OD_600_ equivalent to 1.0 were harvested. A total cell extract was collected by resuspending 0.3 mL of cells with an OD_600_ = 1.0 in 2X Laemmli sample buffer. For the isolation of the periplasmic fraction, cells were fractionated using the EDTA/lysozyme/cold osmotic shock method described above (39). The periplasmic fraction was solubilized by adding an equal volume of 2X Laemmli sample buffer prior to SDS-PAGE.

For samples from the luminescence assay, samples were taken prior to the addition of treatment and at the end of the reaction. Transport was stopped by immediately placing the samples on ice. The total spheroplast extract was collected and the samples were centrifuged at 16,000 g for 2 min at 4 °C. The supernatant was collected as the periplasmic-like fraction (P) and solubilized by mixing with 4X Laemmli sample buffer (Bio-Rad). The pellet was resuspended with the same volume of the 1X M9+ medium and solubilized with 4X Laemmli sample buffer. All samples were heated at 80 °C for 3 min and subsequently subjected to SDS-PAGE and immunoblotting.

### SDS-PAGE and Immunoblotting

Samples were subjected to SDS-PAGE. Upon transfer onto PVDF membranes and blocking, an α-FLAG (GenScript) monoclonal antibody was used to probe for FLAG-tagged SufI or FLAG-tagged SufI-pep86 followed by a horseradish peroxidase-conjugated α-mouse antibody (Santa Cruz Biotechnology Inc.). Labeled proteins were visualized using either ProSignal Pico or ProSignal Femto ECL Western Blotting detection kit (Prometheus).

### Data Analysis

Raw NanoLuc luminescence readouts were normalized using RStudio (Ver. 1.4.1103). For calculating the percentage of luminescence, the luminescence reading obtained immediately before treatment addition was set as 100% (i.e., the luminescence reading baseline signal). Relative luminescence was defined by the difference between the raw luminescence signal and the baseline signal. For the calculation of the relative luminescence values, the luminescence reading baseline was calculated by averaging the signal in the first 20 sec. Normalized data sets were imported into GraphPad Prism version 9.3.0 for macOS (GraphPad Software, San Diego, California USA, www.graphpad.com), where graphs were generated and annotated.

## Acknowledgements

We gratefully acknowledge support from the Division of Chemical Sciences, Geosciences, and Biosciences, Office of Basic Energy Sciences of the U.S. Department of Energy through Grant DE-SC0020304 to SMT.

---

**Supplemental Figure 1.**
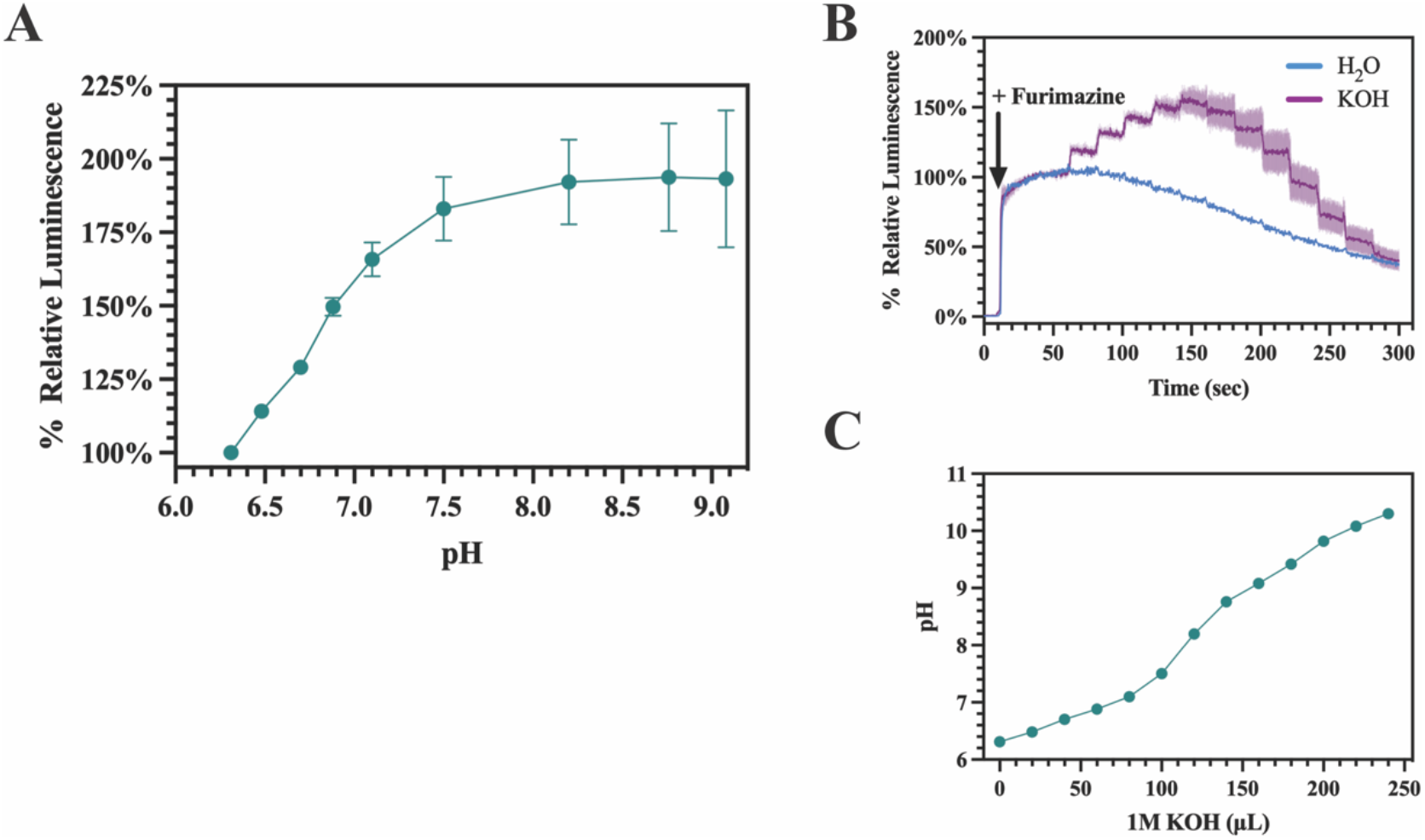
Effects of pH on NanoLuc signals. (**A**) pH dependence of the NanoLuc signal. The assay was performed as in Fig. 4 with isolated SufI-Cpep86 addition to 1X M9^+^ reaction medium containing the 11S protein and furimazine. Error bars represent the standard error from three replicates; luminescence signal from the pH = 6.3 was set as 100%. (**B**) Primary data of the NanoLuc luminescence. 20 μL of 1M KOH or H2O was added to 2 mL of 1x M9^+^ in a stirred cuvette every 20 sec; a pH electrode was also placed in the cuvette. Three replicates for the KOH group were included. Arrow indicates the time when furimazine was added. (**C**) pH titration curve of the 1X M9^+^ reaction medium.

**Supplemental Figure 2.**
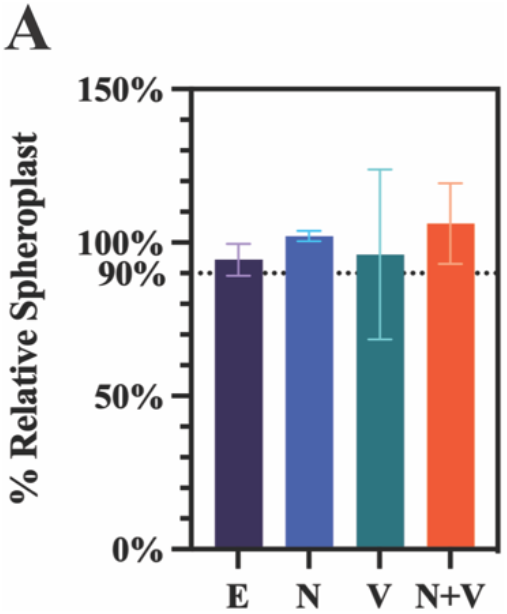
Comparison of the amount of spheroplasts remaining at 50 min post-IPTG induction relative to the amount present at 45 min post-IPTG induction in Figure 6. The spheroplast-to-whole-cell percentages were calculated from the differential lysis of spheroplasts in water as described in Materials and Methods. The bar plot denotes the spheroplast-to-whole-cell percentage at the end of the Fig. 6 experiment (i.e., 50 min post-IPTG induction) relative to the spheroplast-to-whole-cell percentage at the beginning of the Fig. 6 experiment (i.e., 45 min post-IPTG induction); standard error of 3 measurements is shown. E, ethanol. N, 0.5 μM of nigericin. V, 0.5 μM of valinomycin. N+V, 0.5 μM of nigericin and 0.5 μM of valinomycin added together.

